# Chamaeleo: a robust library for DNA storage coding schemes

**DOI:** 10.1101/2020.01.02.892588

**Authors:** Zhi Ping, Haoling Zhang, Shihong Chen, Qianlong Zhuang, Sha Joe Zhu, Yue Shen

## Abstract

Chamaeleo is currently the only collection library that focuses on adapting multiple well-established coding schemes for DNA storage. It provides a tool for researchers to study various coding schemes and apply them in practice. Chamaeleo adheres to the concept of high aggregation and low coupling for software design which will enhance the performance efficiency. Here, we describe the working pipeline of Chamaeleo, and demonstrate its advantages over the implementation of existing single coding schemes. The source code is available at https://github.com/ntpz870817/Chamaeleo, it can be also installed by the command of pip.exe, “pip install chamaeleo”. Alternatively, the wheel file can be downloaded at https://pypi.org/project/Chamaeleo/. Detailed documentation is available at https://chamaeleo.readthedocs.io/en/latest/.

**Author Summary:** DNA is now considered to be a promising candidate media for future digital information storage in order to tackle the global issue of data explosion. Transcoding between binary digital data and quanternary DNA information is one of the most important steps in the whole process of DNA digital storage. Although several coding schemes have been reported, researchers are still investigating better strategies. Moreover, the scripts of these coding schemes use different programming languages, software architectures and optimization contents. Therefore, we here introduce Chamaeleo, a library in which several classical coding schemes are collected, to reconstruct and optimize them. One of the key features of this tool is that we modulize the functions and make it feasible for more customized way of usage. Meanwhile, developers can also incorporate their new algorithms according to the framework expediently. Based on the benchmark tests we conducted, Chamaeleo shows better flexibility and expandability compared to original packages and we hope that it will help the further study and applications in DNA digital storage.

## 1. Introduction

Compared to the orthodox information storage media, DNA has extremely high information capacity and durability [1] and thus is considered to be a future storage medium with great potential. As DNA synthesis and sequencing technology developing, the end-to-end workflow for DNA-based data storage has been well established [2, 3]. Nevertheless, current technology still has particular constrains on DNA sequences such as GC content, length of homopolymer, etc. [4, 5] Therefore, it is a vital issue that how to implement bit-base transcoding with high coding density while fulfilling the biochemical constrains on sequences.

Since 2012, many scientists devoted to developing efficient strategies to store digital information in DNA. Several well-established coding schemes have been proposed in recent years [2]. These schemes focus on optimizing the coding efficiency under certain constrains. Simple code [1], Goldman’s code [6], and Grass’ code [7] use different algorithms to increase coding efficiency as well as eliminate homopolymer. DNA Fountain [8] scheme, which is based on LT code, improves the coding efficiency and applies screening steps to restrict the occurrence of homopolymer and limit GC content. Yin-Yang code [9] provides 1536 derived rules by incorporation of Yin and Yang rules and can be used to find the most suitable rule for a particular file based on its byte frequency [10].

Although programs had been provided for each scheme in their report, their programming languages, software architectures and optimization contents are different, which bring difficulties for subsequent new investigators to start their research in this field. In addition, it is not convenient to compare the performance of different schemes under identical condition or to achieve customized application under specific requirements. Meanwhile, these individual programs do not have well-designed software architecture [11] from the perspective of software engineering. A few program-based validation and improved work will be hindered.

Further studies of DNA-based data storage on transcoding algorithms, data processing and other aspects are still on going. Hence, it is necessary to bring the well-established coding schemes together with fine software structure and provide flexible ways of their usage for research, application, and optimization. Through this work, we introduce Chamaeleo, an open-source library focusing on different coding schemes for DNA storage. This library provides a useful platform for researchers to study different coding schemes. Its efficient implementation is suitable for supporting real-world DNA storage applications.

## 2. Design and Implementation

### 2.1. Library Overview

Currently, Chamaeleo provides five classical coding schemes, including Simple code, Goldman’s code, Grass’ code, DNA Fountain code, and Yin-Yang code. In addition, Chamaeleo also provides independent modules for data process, customized message output, and error correction. We make some minor optimization on original coding schemes for the purpose of better implementation. The details and library tutorials are shown in **S1_Text**.

Chamaeleo has several features:

- Readability: To improve the readability for developers afterwards, this library provides a naming convention in which the identifiers can denote their types and functions.
- Flexibility: We encapsulate [12] different coding schemes and transcoding pipelines. Flexible usages of various hyper-parameters in coding schemes, index operation and error-correction operation are also allowed. Users can further customize their requirements based on this feature.
- Maintainability and Expandability: The library contains five modules, including transcoding module, data handle module, process monitoring module, optional error-correction module, and customized message output module. The modularization of workflow, compared to individual implementations of particular coding schemes, can coordinate arbitrarily and help users to customize the encoding and decoding process flexibly.

Chamaeleo can be used for the encoding and decoding process for DNA storage in the terminal and visual interface such as PyCharm. In **Fig 1**, we demonstrate how Chamaeleo can be used to complete the transcoding process. Specifically, it is divided into two main processes: encoding process and decoding process.

**Fig 1.**
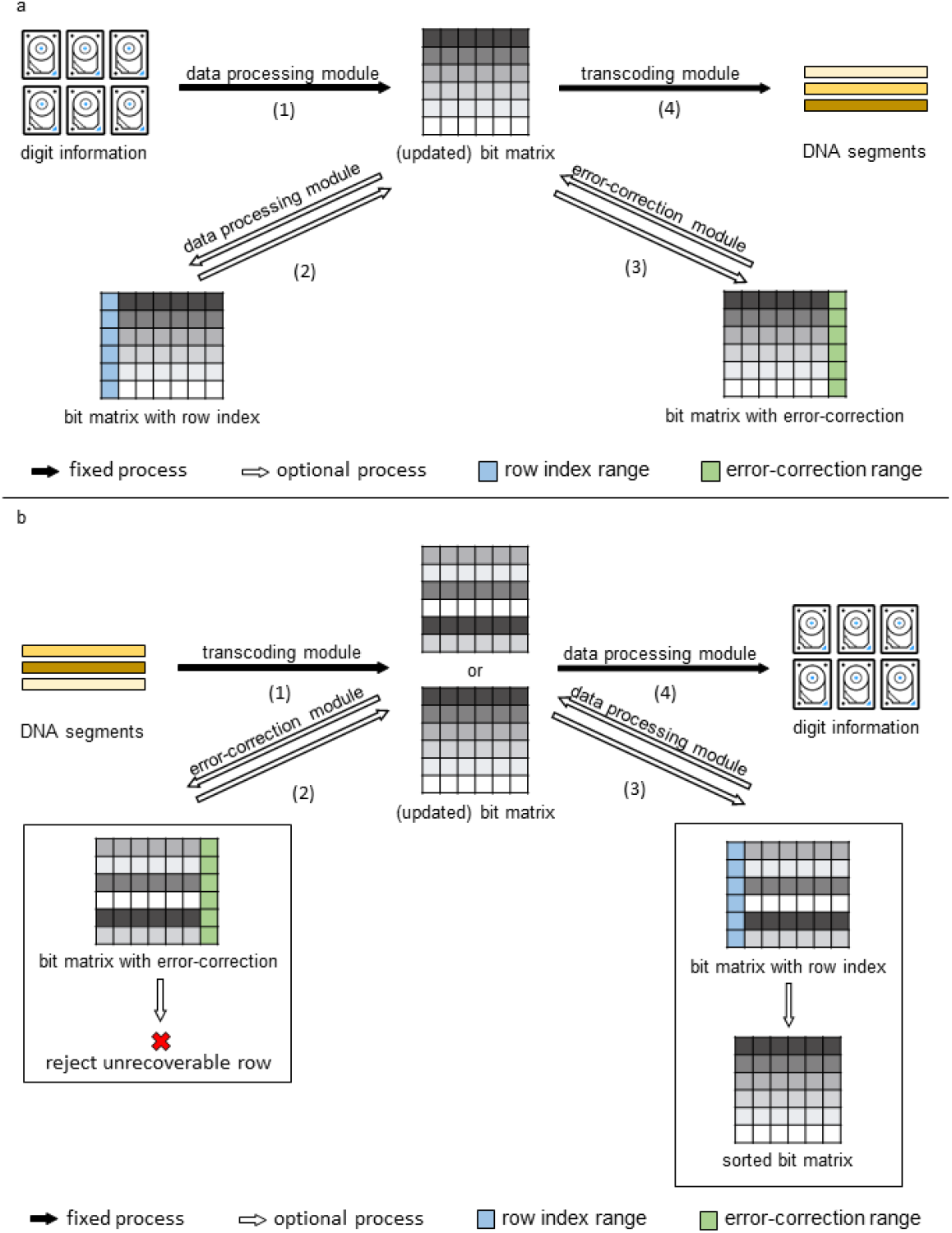
Detailed Illustration of Encoding and Decoding Processing. Step 1 and 4 in (a) and (b) are required, and step 2 and 3 are optional. The complete execution process considering all options is: (1)->(2)->(3)->(4). In the decoding process, if one row is wrong and unrecoverable, the program will reject this row and continue to verify other rows. The processing with flowchart style is shown in **S1_Fig**.

### 2.2. Comparison to Current Individual Programs

Chamaeleo not only integrates the previous well-established coding schemes, but also makes them modulization [13]. The comparison between Chamaeleo and other existing individual programs in the programming contents is shown in **Table 1**.

**Table 1.** Comparison the Programming Contents of Each Tool or Library. √ means the included module in the corresponding tools. The detailed comparisons and evaluations is shown in **S2_Text**.

| Programming Contents | Repositories |  |  |  |  |  |
| --- | --- | --- | --- | --- | --- | --- |
|  | Simple Code | Goldman's Code | Grass' Code | DNA Fountain Code | Yin-Yang Code | Chamaeleo |
| Simple Code | √ |  |  |  |  | √ |
| Goldman's Code |  | √ |  |  |  | √ |
| Grass' Code |  |  | √ |  |  | √ |
| DNA Fountain Code |  |  |  | √ |  | √ |
| Yin-Yang Code |  |  |  |  | √ | √ |
| Parity-check Code |  | √ |  |  |  |  |
| Hamming Code |  |  |  |  | √ | √ |
| Reed-Solomon Code |  |  |  | √ | √ | √ |
| Data Handle Module |  |  |  | √ | √ | √ |
| Customized Message Module |  |  |  |  | √ | √ |

All well-established coding schemes are integrated in Chamaeleo. The encoding and decoding processes of each scheme, with the initial method, are implemented as an individual class in the software architecture [14]. According to our previous study [9], we added some warning prompts in the class of DNA Fountain Code [8], to remind users that with inputted parameters, encoding or decoding process may take very a long time or even fail. In error-correction module, classical Hamming Code and Reed-Solomon code are also implemented as individual classes. Classes in data handle and customized message module are categorized as process classes [14].

### 2.3. Encoding Process

The illustration of encoding process is shown in **Fig 1(a)**. A digit file specified by users is first converted to bit matrix by data process module. The process of adding row indexes with bit format is optional depending on codes. Error-correction can then be added in the above binary matrix optionally by error-correction module. Currently, Hamming code [15] and Reed-Solomon code [16] were included. This library converts the matrix into DNA segments by using the code initialized by users in the transcoding module, and the file consist of DNA segments is finally outputted through data process module.

### 2.4. Decoding Process

This process and the encoding process are complementary to each other, which is shown in **Fig 1(b)**. When the error-correction module was added in their customized process, the incorrect and unrepairable data will be discarded and user will be notified. Later, the matrix will be sorted by its row indexes in this process when the user added the row index in the encoding process.

### 2.5. Different Modes of Usage

The library can be used in two modes: basic and customized. Basic mode is used to implement the transcoding process with default basic parameters preset for each coding scheme. For another, customized mode allows users to modify the parameters such as length of output sequence, desired GC content interval, etc., for customized application.

The basic mode is using the transcoding process once. In addition, users can also consider using the error-correction module once and also adding row indices before each bit segment once. This statement can by executed by visual interfaces or command line.

Considering the many parameters in this statement, we recommend using the templates to accomplish these required tasks. In this situation, it is feasible to use the execution statement of codec_factory.py (in Chamaeleo) directly.

When the highly customized encoding and decoding programs for DNA storage are needed, the basic functions of codec_factory.py can provide will not be able to meet. We strongly recommend using the Python programming tools to accomplish these required tasks. All hyper-parameters in the transcoding module and error-correction module can be adjusted, as long as the value of adjusted hyper-parameters meets the requirements of the corresponding method. In addition, users also can insert their created transcoding method that meets the requirements of the transcoding module, for completing the transcoding process. Before attempting a highly customized transcoding process, the detailed tutorial of Chamaeleo (see **S1_Text**) needs to read.

## 3. Results

### 3.1. A Demo Case in Chamaeleo

In one of our integration testing, we demonstrate the transcoding reliability and correctness in Yin-Yang Code (under random seed **0**) with the file “**Mona Lisa.jpg**”, using codec_factory.py, as a demo case. The hyper-parameters are: (1) rule index **495**, support bases **A**, max ratio **0.8**, search count **10**; (2) need index **True**, verify **Hm()** (in **Chamaeleo/methods/verifies/**), and target DNA segment length **150**.

In the encoding process, the transforms and trends of data are as followed:

1. The file “**Mona Lisa.jpg**” is read by **data_handle.py** (in **Chamaeleo/utils**) for obtaining the original bit matrix.
2. Based on the row (index) length of the obtained bit matrix, the maximum length of index is calculated. The index matrix of the row indices with bit format is connected to the left of the original bit matrix. These operations take place in **index_operator.py** (in **Chamaeleo/methods/components/**).
3. The error-correction bit matrix is created by **Hm.py** and each row of the matrix generated in the last step (M*_l_*). The error-correction bit matrix is connected to the right of M*_l_*, to be the final bit matrix (M*_f_*).
4. DNA sequence set is generated from M*_f_* though **yyc.py** (in **Chamaeleo/methods/**). The detailed transcoding process is shown in [9].
5. The file named “**Mona Lisa.jpg**” is writted by **data_handle.py**, which contains all the generated DNA sequences.

“**Mona Lisa.dna**” becomes “**Mona Lisa.jpg**” through the following 5 steps:

1. The file “**Mona Lisa.dna**” is read by **data_handle.py** for obtaining the whole DNA sequences.
2. The original bit matrix is converted from the above DNA sequences by **yyc.py**.
3. The original bit matrix is verified by **Hm.py**. If the ith line has mistake and cannot be repaired, delete this line and report users. On the Contrary, a novel matrix is obtained by deleting the matrix with error-correcting designated part.
4. Through **index_operator.py** to restore row order in the above matrix and delete the row index part in this matrix (as final bit matrix).
5. Using **data_handle.py**, this final matrix is transformed to “**Mona Lisa.jpg**”.

### 3.2. Benchmarking

In this work, we benchmark the transcoding module and the data process module, respectively. All tests were performed in the environment of Ubuntu 14.04 and Python 3.7.3, with CPU of Intel Core i5 7th Gen.

#### 3.2.1 For the Transcoding Module

Different structures of data can affect the runtime. Therefore, ten files in S1_File with different attributes were collected for benchmarking, including text, picture, audio, video, and typical executive files. Many files were used to perform encoding and decoding processes according to different coding schemes and the runtime was recorded.

As shown in **Table 2**, the file size and the encoding runtime, using identical coding scheme generally exhibits a linear relationship. The encoding runtime of different methods describes the actual time complexity. Because Goldman’s Code and Grass’ Code are encoded with several bits one time, their encoding speeds are faster than other coding schemes. However, the transcoding runtime using some methods may not follow this relationship because of the byte frequency in some files. The extra validity screening process of Yin-Yang Code needs more time, hence longer time is requested for encoding the file “**exiting the Factory.flv**” by Yin-Yang code.

**Table 2.** Encoding Runtimes of Coding Schemes Operating Different Types of File. The unit of **Size** is **KB**, and the unit of encoding runtime is **second**. The segment length in all tested coding schemes is **150** without error-correction code. The hyper-parameters in all tested coding schemes are default (see Detailed Tutorial **S1_Text**).

| File |  |  | Encoding Runtimes |  |  |  |  |
| --- | --- | --- | --- | --- | --- | --- | --- |
| Name | Format | Size | Simple Code | Goldman's Code | Grass' Code | DNA Fountain Code | Yin-Yang Code |
| Mona Lisa | jpg | 96 | 2.68 | 1.57 | 1.55 | 4.64 | 1.57 |
| Microsoft Winmine | exe | 117 | 3.06 | 1.84 | 1.62 | 5.54 | 2.57 |
| The Wandering Earth | pdf | 368 | 7.89 | 4.45 | 3.78 | 19.20 | 5.78 |
| United Nations Flag | bmp | 469 | 10.24 | 5.59 | 4.87 | 57.66 | 7.57 |
| A Tale of Two Cities | pdf | 1011 | 21.42 | 11.90 | 8.89 | 59.44 | 16.69 |
| DNA Fountain Input Files | tar | 2096 | 43.16 | 25.36 | 20.38 | 127.24 | 33.52 |
| For Elise | wma | 2544 | 52.81 | 28.47 | 25.43 | 195.89 | 41.54 |
| I Have a Dream | mp4 | 3986 | 83.14 | 43.26 | 36.08 | 272.61 | 70.39 |
| Exiting the Factory | flv | 4033 | 84.93 | 44.31 | 36.86 | 302.64 | 152.42 |
| Summer | mp3 | 6033 | 126.28 | 75.77 | 59.70 | 422.59 | 107.25 |

As shown in **Table 3**, the decoding speed of Simple Code, Grass’ Code, and Yin-Yang Code are faster than their corresponding encoding speed. There is no big difference in decoding runtime between these three coding schemes. In addition, because the time complexity of Huffman tree decoding process is 5 to 6 times of its encoding process, Goldman’s Code will spend more time to decode DNA sequences than the above three coding schemes.

**Table 3.** Decoding Runtimes of Coding Schemes Operating Different Types of File. The units of table and the hyper-parameters of all tested coding schemes are same as the **Table 2**. “error-1” is caused by the decoded DNA sequences is not full rank [17] and “error-2” is caused by the system crash from the excessive recursive calls in Python [18].

| File |  |  | Decoding Runtimes |  |  |  |  |
| --- | --- | --- | --- | --- | --- | --- | --- |
| Name | Format | Size | Simple Code | Goldman's Code | Grass' Code | DNA Fountain Code | Yin-Yang Code |
| Mona Lisa | jpg | 96 | 0.52 | 9.98 | 0.75 | 3.19 | 0.90 |
| Microsoft Winmine | exe | 117 | 0.56 | 12.53 | 1.21 | 5.30 | 1.13 |
| The Wandering Earth | pdf | 368 | 1.88 | 38.91 | 2.36 | 12.80 | 2.22 |
| United Nations Flag | bmp | 469 | 2.00 | 49.28 | 3.15 | error-1 | 2.67 |
| A Tale of Two Cities | pdf | 1011 | 5.81 | 66.53 | 7.89 | 127.79 | 6.05 |
| DNA Fountain Input Files | tar | 2096 | 11.04 | 237.30 | 14.95 | 528.72 | 13.71 |
| For Elise | wma | 2544 | 14.70 | 283.35 | 18.85 | error-2 | 16.61 |
| I Have a Dream | mp4 | 3986 | 21.67 | 461.96 | 30.34 | error-2 | 26.62 |
| Exiting the Factory | flv | 4033 | 22.40 | 420.10 | 31.59 | error-2 | 28.52 |
| Summer | mp3 | 6033 | 34.40 | 683.58 | 48.79 | error-2 | 39.98 |

#### 3.2.2. For the Data Process Module

The byte frequency does not affect the runtime of the data process module. So we use the bit data consisting of “**0**” and DNA segment consisting of “**A**” in this benchmark test. After 100 sets of data testing, the runtime of transcoding are as follows: 4.45 Mebibyte/s (for reading binary), 1.84 Mebibyte/s (for writing binary), 0.72 Mebibase/s (for reading DNA), and 0.21 Mebibase/s (for writing DNA).

In the default setting of Chamaeleo, bit segment (payload) length is set as 120 to allow users add flanking sequences based on the length limitation of common industrial oligo synthesis, 200nt to date. The information density of DNA-based data storage is determined not only by bit-base transcoding algorithm itself but by the index length in each output sequence as well. With fixed output sequence length and increasing file size, the index length will also increase and thus reduce the length of data region. Therefore, to maintain a shorter row index length is an important issue. For example, when the size of file reaches 15GB, the length of row index will reach 30 bits and the coding efficiency will be reduced to 80% of the original. Two strategies are recommended when large file is transcoded: (1) increase the bit segment length if users are able to synthesize longer oligonucleotide; (2) divide the file to more parts for storage.

In addition, the decoding runtime and decodability of coding scheme also need to be evaluated by users. In DNA Fountain Code, we recommend users not to transcode digital files more than 3MB, because of overlong decoding runtime and overlarge computation.

#### 3.3. Error Model

Sometimes a DNA segment cannot be correctly retrieved to bit segment(s) in the decoding process. Some operations, such as DNA synthesis and sequencing, may cause errors in DNA segments including insertion, deletion or mutation. Here, we demonstrated the impact of these errors on the decoding process of the above-mentioned coding schemes.

Three main strategies are usually used to correct errors in DNA storage: (1) Increasing logical redundancy [19]; (2) Adding error-correction; (3) Self-checking in the coding scheme.

However, these strategies fail to resolve errors caused by insertion and deletion. In practice, if these two kinds of errors are encountered, such DNA segments usually be discarded. Therefore, we only discuss the case of mutation errors here.

#### 3.3.1. Increasing Correct Backup by Redundancy

Additional redundancy helps to increase the recovery stability from DNA sequences to digital files [2] and reduce the probability of systematic failure. There are two ways to increase redundancy [20] (see **Fig 2**): one is splitting the inputted DNA sequences into overlapping segments to provide fourfold redundancy for each segment, the other is taking the exclusive-or of two DNA segment to form a third. Two kinds of redundancy are remarkable in the coding schemes.

**Fig 2.**
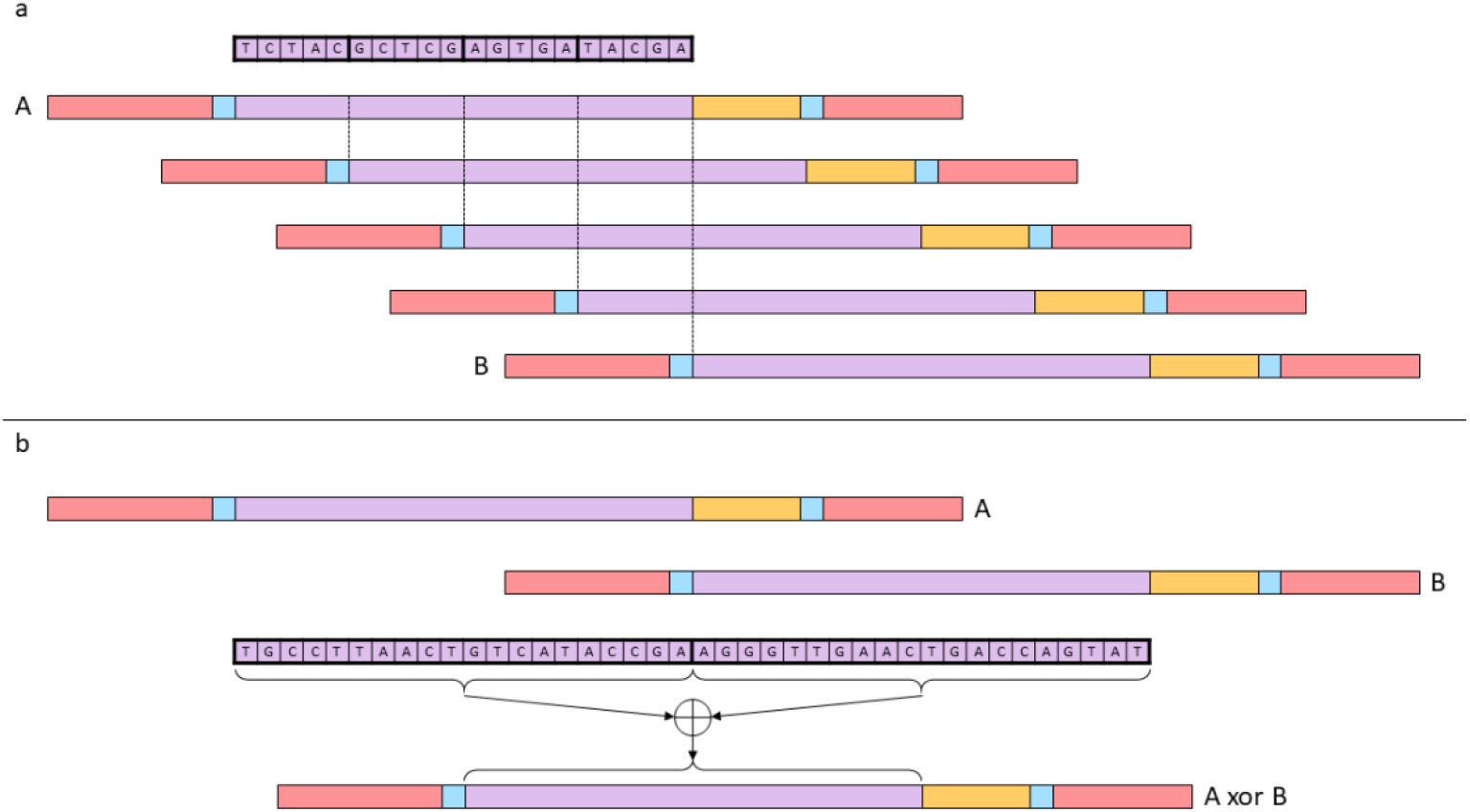
Two Ways to Increase Redundancy in DNA Storage. In panel a [6], each DNA segment is split into four overlapping fragments. Between A and B sequences, three additional sequences are generated in turn, thus, the same segment is magnified four times. In panel b [20], any two of the three segments A, B, and (A xor B) are sufficient to recover the three.

When these DNA segments have enough redundant DNA segments for verification, they can be decoded completely correctly. However, adding redundancy will reduce the coding efficiency of the coding scheme itself. For the first way, see **Fig 2(a)**, when the redundancy is *k* times, the coding efficiency of the coding scheme will becomes 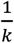 of the original one. For the other way, see **Fig 2(b)**, the coding efficiency of the coding scheme will becomes 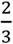 of the original one. The theoretical density of this way is much higher than the first way.

#### 3.3.2. Error-correction Strategy for Restoring the Error Information

Based on experimental data, we get three kinds of error possibilities: insertion = 0.075%, deletion = 0.075%, and mutation = 0.15%.

We set *m* for the length of bit segment and *s* for the number of additional check bits. The code rate of Hamming Code and Reed-Solomon Code are 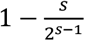 and 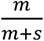 respectively. And their error-correction capability are 1 and 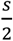 in turn. The code rate also describes the coding efficiency. The product of code rate and coding efficiency of the coding scheme determines the final coding efficiency of the encoding process. When additional check bits become larger, the coding efficiency of the encoding process decreases significantly. Therefore, there is a negative correlation between the error-correction capability and the coding efficiency of the encoding process. Interestingly, based on the error possibilities of short segments is very low (maybe the whole DNA segment has only one mutation error), the use of error-correction strategy is a more economical solution than that of redundancy strategy.

#### 3.3.3. Self-checking by Restrictions of Coding Scheme

Based on the constraints of the coding scheme, some coding schemes have self-checking function. They can pre-verify whether current DNA segments can be decoded, even repair the incorrect DNA segment. For example, if there are two consecutive identical bases in the DNA segment obtained by Goldman’s code, this DNA segment must be problematic. So we count the detectability of all the above coding schemes, which are shown in **Table 4**.

**Table 4.** Self-checking Behavior of Coding Schemes. The detection rate is the expected frequency that the coding scheme itself can find the error in the corresponding DNA sequences. The detection bias represents the possible wrong location range when the coding scheme meets the error. nv(*m, n*) describes the rate of DNA segment with *m* length no longer satisfy constraints after mutating *n* bases. If the constraints does not exist, all the mutating situations meet the constraints, the extreme value of nv(*m, n*) is 0, as same as Simple Code.

| Parameters | Coding Schemes |  |  |  |  |
| --- | --- | --- | --- | --- | --- |
|  | Simple Code | Goldman's Code | Grass' Code | DNA Fountain Code | Yin-Yang Code |
| Detection Rate | 0 | $\frac{\binom{1}{4}}{\binom{1}{4} \times \binom{1}{4}} = \frac{1}{4}$ | $1 - \frac{\binom{1}{4} \times \binom{1}{4} \times \binom{1}{3} - 1}{\binom{1}{4}^3} = \frac{17}{64}$ | $\sum_{n=1}^m nv(m, n)$ | |
| Detection Bias | N/A | 2 | 3 | depends on constraints |  |

These coding schemes have a certain probability of detecting errors and obtaining the interval of the error location. However, among the known coding schemes have no real sense of self-checking. Because it does not accurately locate errors, let alone repair these errors. We look forward to a coding scheme that can detect errors and fix them in the future.

## Discussion

Files with different attributes were transcoded into DNA sequences using incorporated coding schemes. The runtime of encoding and decoding using different coding schemes were benchmarked. The results were proved to be consistent with the comparison between these schemes using previous packages. Furthermore, the unique data handle module introduced in this library, handling bit arrays and DNA sequences, also showed a fast performance for reading and writing of both binary and DNA sequences. We anticipate that this library will facilitate the development of future tools of DNA storage and help to create more a comparative model for different coding schemes.

This library would be an open source tool for developers and users. In the future, we expect more developers to incorporate their unique coding algorithms or strategies for DNA digital storage into this library in the future. Two issues can be tackled in the next version: (1) incorporate more coding schemes as well as their unique module, like Composite DNA letters [21]; (2) establish an evaluation system and criteria which can compare different coding schemes.

## Supporting Information

### S1 Text

**Detailed Tutorial of Chamaeleo.** This document describes the general and customized usage of programming code. (PDF)

### S2 Text

**Detailed Comparison between Chamaeleo and other individual programs.** (PDF)

### S1 Fig

**Workflow of the Chamaeleo.** “Encoding process” from binary code to DNA segment, and “Decoding process” from DNA code segment to binary code.

### S1 File

**Ten Files in the Benchmarking.** (ZIP)

## Acknowledgments

This work was supported by National Key Research and Development Program of China (No. 2018YFA0900100), the Guangdong Provincial Key Laboratory of Genome Read and Write (No.2017B030301011) and China National GeneBank, Shenzhen.

## Author Contributions

Software: *HLZ*, *SJZ* and *ZP*.

Validation: *HLZ*, *SJZ*, *SHC*, and *QLZ*.

Visualization: *ZP*, *SJZ*, and *HLZ*.

Funding acquisition: *ZP*.

Supervision: *YS*.

Writing in original draft: *ZP* and *HLZ*.

Writing in review & editing: *ZP*, *HLZ*, *SJZ*, *SHC*, and *YS*.

