## Supplementary material for "Chamaeleo: a robust library for DNA storage coding schemes": S1 Text

### Detailed Tutorial of Chamaeleo

#### Basic transcoding requirement

The basic transcoding requirement is using the transcoding process once. In addition, users can also consider using the error-correction module once and also adding row indices before each bit segment once. In this situation, it is feasible to use **codec_factory.py** (in Chamaeleo/) directly.

This statement can by executed by visual interfaces or command line. Considering the many parameters in this statement, we recommend using the templates to accomplish these required tasks. Taking Yin-Yang code (Ping, et al., 2019) as an Example, the specific usage is as followed:

Using Python programming tools:

```
# encode part
from chamaeleo import *
method = yyc.YYC()
codec_factory.encode(method, input_path="C:/init.mp4", output_path="C:/target.dna", model_path="C:/yyc.pkl")
# decode part
codec_factory.decode(method, input_path="C:/target.dna", output_path="C:/target.mp4")
# codec_factory.decode(input_path="C:/target.dna", output_path="C:/target.mp4", model_path="C:/yyc.pkl")
```

Using command line:

```
$ encode part $
> codec_factory.encode(yyc.YYC(), input_path="C:/init.mp4", output_path="C:/target.dna", model_path="C:/yyc.pkl")
$ decode part $
> codec_factory.decode(yyc.YYC(), input_path="C:/target.dna", output_path="C:/target.mp4")
$ > codec_factory.decode(input_path="C:/target.dna", output_path="C:/target.mp4", model_path="C:/yyc.pkl") $
```

In the two statements in **codec_factory.py**, the hyper parameters of encoding and encoding must correspond one by one. In the encoding process, **codec_factory.encode(…)**, the hyper parameters are: **method**, **input_path**, **output_path**, **model_path**, **verify**, **need_index**, and **segment_length**. The decoding process, **codec_factory.decode(…)**, is consistent with the hyper parameters of the encoding process. If **need_index** is **True** in encoding process, **has_index** needs **True** in decoding process. If **verify** is an instantiated method (**=code**) in the encoding process, **verify=code** needs in decoding process.

The default value of hyper parameters are:

| Parameter name | Default value | Description |
|---|---|---|
| **model_path** | None | No use of saving the serialized file of the method. |
| **verify** | None | No use of using error-correction coding module. |
| **need_index** | True | The use of adding row index for each bit segment. |
| **segment_length** | 120 | The default segment length is 120 bits. |

In addition, there are different requirements for the instantiation of hyper parameters in different transcoding methods (in the transcoding module, Chamaeleo/methods/). It will perform hyper parameters checking for each transcoding method before the encoding or decoding program runs. This part is discussed in Section 1.4 in conjunction with highly customized issues.

#### Transcoding requirements with highly customized

When we need highly customized coding and decoding programs for DNA storage, the basic functions of **codec_factory.py** can provide will not be able to meet. We recommend using the Python programming tools to accomplish these required tasks.

In the transcoding module, four transcoding method is collected. Here, we will discuss the hyper parameters of these above four methods respectively.

Simple Code (see **sc.py**, in Chamaeleo/methods) has only one hyper parameter: **mapping_rule**, which describes the mapping between bases and numbers.

```
# initialize Simple Code
tool = sc.SC(mapping_rule=[0, 1, 2, 3])
# tool = sc.SC(mapping_rule=[0, 0, 1, 1])
```

Each position of **mapping_rule** is mapping to 0-A, 1-C, 2-G, 3-T. There can be two settings of this hyper parameter: (1) two nucleotides correspond to a number (0 or 1): i.e. A/T-0, C/G-1; (2) each nucleotide corresponds to a number: i.e. A-00, T-01, C-10, G-11.

Goldman's Code (see **hc.py**, in Chamaeleo/methods) has also one hyper parameter: **fixed_huffman**, which describes whether user uses the fixed Huffman tree or not.

```
# initialize Goldman's Code
tool = hc.HC(fixed_huffman=True)
```

**fixed_huffman** is a Boolean value. The fixed Huffman tree is the mapping between binary and ternary created by Goldman's (see **inherent.py**, in Chamaeleo/methods/components). It prevents some users from losing the Huffman tree of the file itself during decoding process. However, the coding efficiency cannot be effectively guaranteed for arbitrary digital files.

Grass' Code (see **gc.py**, in Chamaeleo/methods) has one hyper parameter: **base_values**, which describes the mapping of bases and 47$^{th}$ digit. Taking the incremental list as an example:

```
# initialize Grass' Code
tool = gc.GC(base_values=[index for index in range(48)])
```

**base_values** is a list of 0 to 47 (total of 48 numbers), which is a one-to-one mapping of 47 groups of 3 nucleotides and 47$^{th}$ digit. Different mappings determine the diversity of DNA sequences produced.

Fountain Code (see **fc.py**, in Chamaeleo/methods), implementing from DNA Fountain, has 6 hyper-parameters: **homopolymer**, **gc_content**, **redundancy**, **c_dist**, **delta**, **header_size**, and **decode_packets**. Here are the descriptions and default values of these hyper parameters.

| Parameter name | Default value | Description |
|---|---|---|
| **homopolymer** | 4 | the maximum length of homopolymer. |
| **gc_content** | 0.2 | the fraction of gc content above/below 0.5 (0.1 means 0.4-0.6). |
| **redundancy** | 0.5 | artificial redundancy for decode successfully (0.5 generate 50% more fragments). |
| **c_dist** | 0.1 | degree distribution tuning parameter. |
| **delta** | 0.5 | degree distribution tuning parameter. |

| | | |
|---|---|---|
| **recursion_depth** | 10000000 | adjust the maximum recursion depth in Python. |
| **header_size** | 4 | number of bytes for the header. |
| **decode_packets** | None | bit segments in the encoding process. If it is missing during decoding process, you need to declare it when initializing the coding scheme. |

For the default requirements of Fountain Code, the specific usage is as followed:

```
# initialize Yin-Yang Code
tool = fc.FC()
# tool = yyc.YYC(homopolymer=4, gc_content=0.2, redundancy=0.5, c_dist=0.1, delta=0.5,
recursion_depth=10000000, header_size=4, decode_packets=None)
```

There are many hyper parameters of Yin-Yang Code, **yyc.py** (in Chamaeleo/methods), which makes it have at least 6,144 different mapping modes. Its hyper parameters are: **base_reference**, **current_code_matrix**, **support_bases**, **support_spacing**, **max_ratio**, and **search_count**. Here are the descriptions and default values of these hyper parameters.

| Parameter name | Default value | Description |
|---|---|---|
| **base_reference** | [0, 1, 0, 1] | Yang rule, correspondence between base and bit data in the upper bit segment. The default value is Rule 495. |
| **current_code_matrix** | [[1, 1, 0, 0], [1, 0, 0, 1], [1, 1, 0, 0], [1, 1, 0, 0]] | Yin rule, correspondence between base and bit data in the lower bit segment. The default value is Rule 495. |
| **support_bases** | A | Base replenishment before official data. |
| **support_spacing** | 0 | Spacing between support base and current base. When the support base is the front of the current base, the spacing is 0. |
| **max_ratio** | 0.8 | The max ratio of 0 or 1. When the (count/length) >= this parameter, we decide that this binary sequence is not good. |
| **search_count** | 1 | Maximum number of queries. If the DNA segment generated by two bit segments obtained in the current query is valid, the query is stopped. |

For the default requirements of Yin-Yang Code, the specific usage is as followed:

```
# initialize Yin-Yang Code
tool = yyc.YYC()
# tool = yyc.YYC(base_reference=[0,1,0,1], current_code_matrix=[[1,1,0,0],[1,0,0,1],[1,1,0,0],[1,1,0,0]],
support_bases=‘A’, support_spacing=0, max_ratio=0.8, search_count=1)
```

#### Error-correction method customization

If the transcoding process needs to be highly customized, like add error-correction code repeatedly, **codec_factory.py** will not be able to solve the demand. This module, Hamming code (**hm.py** in Chamaeleo/methods/verifies) and Reed-Solomon code (**rs.py** in Chamaeleo/methods/verifies), can deal with one-dimensional and two-dimensional data. Considering that parity check has no correction function, we do not put it into this library. But we have interfaces for it.

Each code has three operations: add, remove, and verify. Among them, verify operation has repair function, which can fix the wrong data within a certain limit (see Section 6.2). Here, we take Hamming code as an example:

```
# initialize Hamming code
ec = hm.Hm()
# add Hamming code
input_matrix = ec.add_for_matrix(input_matrix)
# verify Hamming code
output_matrix = ec.verify_for_matrix(output_matrix)
# remove Hamming code
output_matrix = ec.remove_for_matrix(output_matrix)
```

If the users need to reuse Hamming codes, the specific usage is as followed:

```
# add Hamming code 10 times
for i in range(10)
    input_matrix = ec.add_for_matrix(input_matrix)
```
