## Supplementary material for "Chamaeleo: a robust library for DNA storage coding schemes": S2 Text

### Detailed Comparison Between Programs

#### Simple Code

| | |
|---|---|
| **Source** | Next-generation digital information storage in DNA |
| **Last update** | 17 Aug 2012 |
| **Links** | https://science.sciencemag.org/content/suppl/2012/08/15/science.1226355.DC1 |
| **Functions** | ● Encode by simple code.<br>● Decode by simple code. |
| **Difference** | Chamaeleo contains all the above functions and integrates the transformation part into a class. In addition, it provides two initialization strategies for the transcoding rule. |
| **Comments** | Simple Code is implemented by Perl and Python without modularization. |

#### Goldman's Code

| | |
|---|---|
| **Source** | Towards practical, high-capacity, low-maintenance information storage in synthesized DNA |
| **Last update** | 5 Mar 2018 |
| **Links** | https://www.ebi.ac.uk/goldman-srv/DNA-storage/ |
| **Functions** | ● Encode by Goldman's code.<br>● Decode by Goldman's code.<br>● Find errors by Parity-check code. |
| **Difference** | Chamaeleo contains all the above functions and integrates the transformation part into a class. In addition, it provides two initialization strategies for the Huffman tree: one is using the fixed Huffman tree (Knuth, 1985), which is created by Goldman the other is created by the digital file provided by user. |
| **Comments** | Goldman's Code does not consider adding the row index for each segment. In Chamaeleo, In Chamaeleo, because the row indices are considered, the length of each DNA segment is not necessarily equal. Sometimes, these DNA segments will be one or two base discrepancies in practical use. The variable-length data processing causes the unstable decoding runtime of Goldman's Code. |

#### Grass' Code

| | |
|---|---|
| **Source** | Robust chemical preservation of digital information on DNA in silica with error-correcting codes |
| **Last update** | 3 Nov 2014 |
| **Links** | N/A |
| **Functions** | ● Encode by Grass' code.<br>● Decode by Grass' code.<br>● Correct errors using Reed-Solomon code. |
| **Difference** | Chamaeleo contains all the above methods and integrates the transformation part into a class. |
| **Comments** | Considering the transcoding rule of Grass' Code, the length of each bit segment must be a multiple of 16. |

#### Fountain Code

| | |
|---|---|
| **Source** | DNA Fountain enables a robust and efficient storage architecture |
| **Last update** | 2 Mar 2017 |
| **Links** | https://github.com/TeamErlich/dna-fountain |
| **Functions** | ● Encode by Fountain code.<br>● Decode by Fountain code.<br>● Correct errors using Reed-Solomon code.<br>● Read and write digital files, read and write DNA segments (including fasta and non-fasta) by Data Handle Module.<br>● Print logs directly. |
| **Difference** | Chamaeleo provides an interface for this method. |
| **Comments** | The program of Fountain Code divides the method into different modules to some extent. It also speed up by C language. However, it has some minor errors in it programing. Sometimes, the decoding process cannot decode the produced DNA segments for some unknown reasons. |

#### Yin-Yang Code

| | |
|---|---|
| **Source** | Towards Practical and Robust DNA-based Data Archiving by Codec System Named 'Yin-Yang' |
| **Last update** | 9 Nov 2019 |
| **Links** | https://github.com/ntpz870817/DNA-storage-YYC |
| **Functions** | ● Encode by Yin-Yang code.<br>● Decode by Yin-Yang code.<br>● Correct errors using Hamming code or Reed-Solomon code.<br>● Read and write digital files, read and write DNA segments) by Data Handle Module.<br>● Print logs by Customized Message Module. |
| **Difference** | Chamaeleo provides an interface for this method and integrates the transformation part into a class. |
| **Comments** | Because the program of Yin-Yang Code has been optimized by modularization, its transcoding part is copied into Chamaeleo directly. Yin-Yang Code employs an extra validity screening process to ensure the feasibility of DNA sequences, so its runtime of encoding process is based on code frequency of specific file. |
