## Supplementary figures and images for "Chamaeleo: a robust library for DNA storage coding schemes"

### S1 Fig

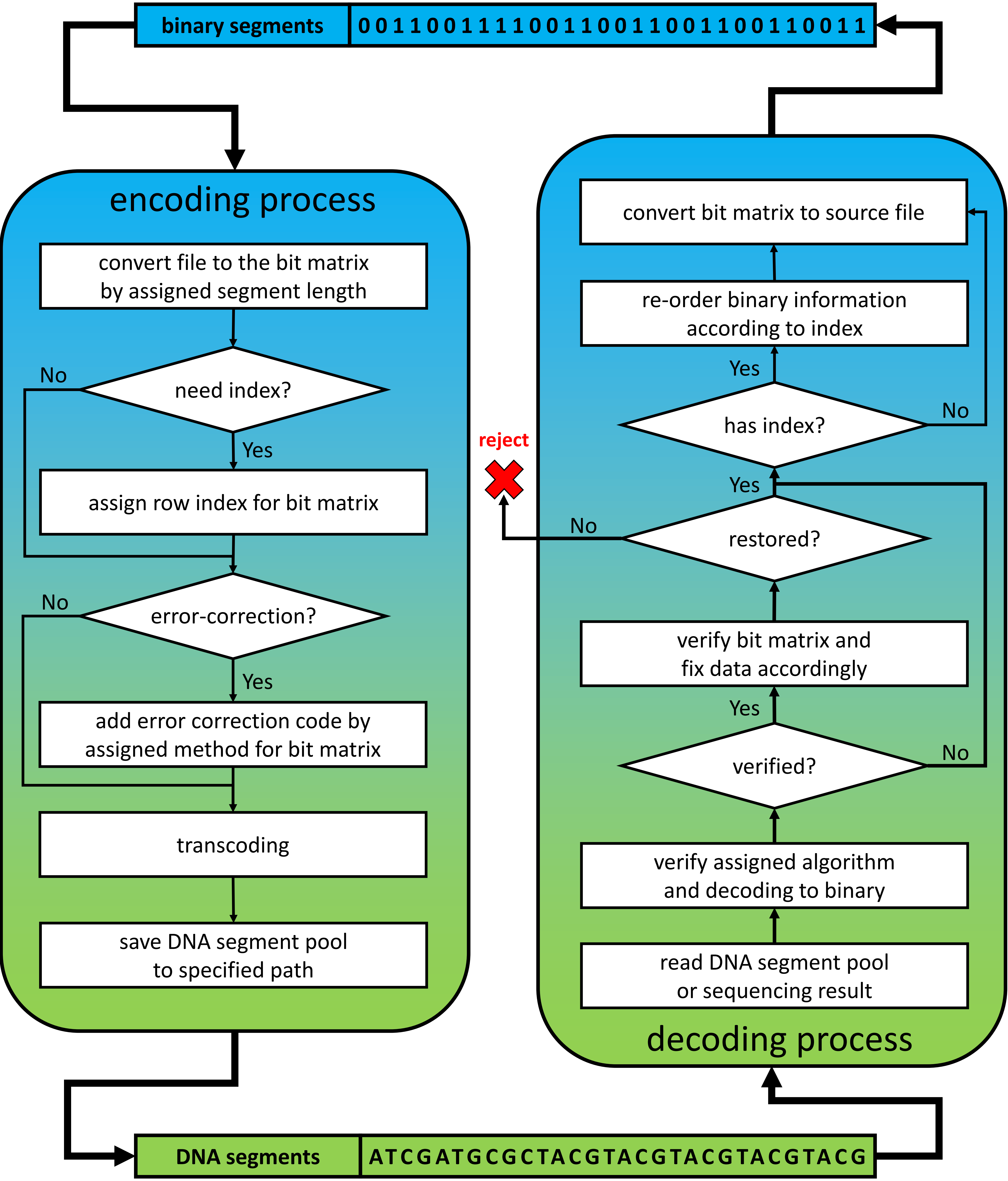
